# Gut microbiota in the burying beetle, *Nicrophorus vespilloides*, provide colonization resistance against larval bacterial pathogens

**DOI:** 10.1101/157511

**Authors:** Yin Wang, Daniel E. Rozen

## Abstract

Carrion beetles, *Nicrophorus vespilloides*, are reared on decomposing vertebrate carrion where larvae are exposed to high-density populations of carcass-derived bacteria. We previously showed that larvae do not become colonized with these bacteria, but instead are colonized with the gut microbiome of their parents. These results suggested that bacteria in the beetle microbiome outcompete the carcass derived species for colonization of the larval gut. Here we test this hypothesis directly and quantify the fitness consequences of colonization of the *Nicrophorus* larval gut with different bacterial symbionts, including the insect pathogen *Serratia marcescens*. First, we show that beetles colonized by their endogenous microbiome produce significantly heavier broods than those colonized with carcass-bacteria. Next, we show that bacteria from the endogenous microbiome, including *Providencia rettgeri* and *Morganella morganii*, are better colonizers of the beetle gut and can outcompete non-endogenous species, including *S. marcescens and Escherichia coli*, during *in vivo* competition. Finally, we find that *Providencia* and *Morganella* provide beetles with colonization resistance against *Serratia* and thereby reduce *Serratia*-induced larval mortality during co-inoculation. Importantly, this effect is eliminated in larvae first colonized by Serratia, suggesting that while competition within the larval gut is strongly determined by priority effects, these effects are less important for Serratia-induced mortality. Our work supports the idea that bacterial gut symbionts provide direct benefits to Nicrophorus larvae by outcompeting potential bacterial pathogens. They further suggest that one benefit of parental care in *Nicrophorus vespilloides* is the social transmission of the microbiome from caring parents to their offspring.

## Introduction

Animals are colonized by a diverse array of bacterial symbionts, the microbiome, that provide essential functions to their hosts (1–3). Animal microbiomes can alter nutrient uptake (4), development (5), parasite susceptibility (6), and even behaviors like mate choice (7). In addition, symbionts that reside within animal guts can provide their hosts with resistance to bacterial pathogens via a process called colonization resistance (8). For example, the gut bacterial community of locusts, *Schistocerca gregaria,* prevents invasion and disease from the insect pathogen *Serratia marcescens,* an outcome that depends in part on the diversity of the gut microbial community (9). Similarly, honeybees became more susceptible to *Serratia* infection following treatment with antibiotics that altered the structure of their endogenous microbiota (10). These results support the idea that gut bacteria can provide protection against pathogens while also highlighting the importance and timing of symbiont transmission in juvenile animals (11). However, it is often unclear if colonization resistance results from specific inhibition of invading pathogens or whether it results from the simple fact that symbionts get there first (9, 12, 13). In other words, is colonization resistance the result of specificity or priority?

To address this question we focus on the role of the endogenous microbiota of the burying beetle, *Nicrophorus vespilloides.* This system is especially well suited to this work given the high exposure of larval beetles to environmental bacteria (14–17), together with extensive data on the composition and transmission of the beetle microbiome from parents to offspring (18–20). *N. vespilloides* larvae are reared on small vertebrate carcasses where they feed directly from the carcass and are provided regurgitated food from parent beetles that care for developing broods (14, 21, 22). Parental beetles dramatically increase larval growth and fitness during brood rearing by investing in pre‑ and post-hatch care (15, 22, 23). During pre-hatch care, parents remove the fur and guts of the carcass and coat its surface in oral and anal secretions that have antimicrobial activity (14, 24–26). Post-hatch, parents defend their developing larvae from other insect species and also feed larvae with regurgitated food (21, 27). In a recent study we found that parents transmit their gut microbiome to their larvae by direct feeding. In addition, we found that the core members of this microbiota could even be transmitted to larvae indirectly, by bacteria deposited onto the carcass by parents (18). This unexpected result suggested that these core bacterial species were outcompeting the numerous microbes living on and inside the carcass within the larval gut, thus giving rise to the stable endogenous *Nicrophorus* microbiome (18, 19). However, the mechanisms of their increased competitiveness remained unclear as were the consequences of their colonization.

Here we carry out invasion experiments into sterile larvae to directly quantify competitive interactions taking place between endogenous and non-endogenous microbes from the *Nicrophorus* gut. We first quantify bacterial growth rates within the larval gut and then directly determine the competitive interactions between species during mixed inoculations and in different orders. Finally, we quantify whether members of the core microbiome provide colonization resistance against *Serratia marcescens,* and known insect pathogen. Briefly, we show that native gut species significantly outcompete foreign species within the host gut, irrespective of infection order. In addition, we find that the endogenous microbiota increases beetle fitness, both in terms of brood size and in terms of pathogen resistance in larvae. Our results provide strong evidence that an important benefit of parental care in *N. vespilloides* is the social transmission of the microbiome from caring parents to their offspring.

## Methods and Materials

### Beetle collection and rearing

Experimental beetles were taken from an outbred laboratory population derived from wild-caught *N. vespilloides* individuals trapped near Leiden in The Netherlands, between May and June 2015. Beetles were maintained in the laboratory at 20°C with a 15:9 hour light: dark cycle and fed fresh chicken liver twice a week. Mating pairs were established by placing a male and female in a small plastic container containing ∼ 1 cm soil overnight. Mated females were provided with a fresh carcass (20-23g) the following morning to initiate egg laying.

To examine the impact of different microbial communities on *N. vespilloides* fitness, we established independent treatment populations containing endogenous or carcass derived gut bacteria, designated FC (full-care) and NC (no-care) beetles respectively. Whereas parents and larvae in the FC treatment were reared in the presence of parental care and thus acquired their microbiota primarily from their parents, larvae in the NC group were reared in the absence of parental care (with an unprepared carcass that we opened using a sterile scalpel), and acquired their microbiota from the carcass and surrounding soil (18). Ten day old adults that had eclosed from FC and NC broods were paired within treatments for mating (n = 15 / treatment) and subsequently provided with a fresh mouse carcass (22-24g) for breeding. The fitness of both parental treatment groups (NC and FC) was determined by quantifying total brood size, total larval weight and mean larval mass.

### Experimental bacterial inoculation of *N. vespilloides* larvae

To generate germ-free larvae, we collected eggs 15 hours after FC females were provided with a fresh carcass. These were surface sterilized twice for 15 minutes in an antimicrobial solution containing hen-egg white lysozyme (1 mg/ml), streptomycin (500 μg/ml) and ampicillin (100 μg/ml), and followed by a sterile water wash. Next, treated eggs were transferred onto 1% water agar plates to hatch. Previous experiments have shown that eggs thus treated are free of bacteria (28). 0 - 24h old first-instar larvae were transferred onto new sterile 1% water agar petri dishes (100mm × 15mm) in groups of a maximum of 7 larvae. Larvae on each plate were derived from independent breeding pairs. Larvae were fed a sterile diet developed using Pasteurized chicken liver prepared via a “Sous vide” cooking approach. Fresh chicken liver was sliced into 3g chunks using aseptic technique and transferred in individual pieces to a 1.5ml eppendorf tube containing 100 μl sterile water. These were then placed in a water bath at 65°C for 8 minutes, followed by immediate cooling at −20°C. We determined the effectiveness of this method by plating liver samples before and after pasteurization onto both 1/3 strength Tryptic Soy agar and LB agar. The initial CFU of unpasteurized liver was ∼ 1e^6^/gram CFU while following treatment the CFU was reduced to 0 (with a limit of detection of ∼ 10 CFU/mL). Larvae were offered this sterile diet, alone or coated with different bacterial inocula, on new 1% water agar plates daily.

### *In vivo* competition within larvae

To determine if “endogenous” bacteria can outcompete foreign strains during larval colonization we competed bacterial strains against one another within the larval gut, focusing on four different bacterial species. The bacterial species *Providencia rettgeri* and *Morganella morganii* are abundant *N. vespilloides* gut symbionts throughout development and are considered “endogenous” species (18, 19). By contrast, *Serratia marcescens* and *Escherichia coli,* which are found commonly in both soil and on decomposing carcasses, colonize larvae that are reared without parental care in NC broods (18). *S. marcescens* is also a known insect pathogen in several insect species (29–31), including *N. vespilloides. P. rettgeri* (P) and *M. morganii* (M) were isolated from *N. vespilloides* adults guts while *S. marcescens* (S) and *E. coli* (E) were isolated from decomposing mouse carcasses (18).

Bacteria for inoculations were cultured overnight at 30°C in 1/3 TSB medium. Overnight cultures of each species were pelleted and washed two times in sterile phosphate buffered saline (PBS, pH= 7.2), and diluted to an optical density at 600nm (OD_600_) of 0.2 measured using a BIO-RAD SmartSpec™ Plus spectrophotometer. Ten microliters of this solution, containing ∼10^6^ cells total, was used to coat sterile liver prepared as above. Inoculations with 2 species contained the same total bacterial density, with each species present at a 1:1 ratio. Larvae were provided with inoculated diet for six hours on a sterile water agar plate, after which they were transferred to a new agar plate containing new sterile diet. Subsequent transfers to plates containing fresh sterile food took place every 24 hours for 7 days, or until larvae were destructively sampled. In experiments where larvae were sequentially challenged with different bacterial species, we treated larvae the same as above, but larvae were inoculated with target strains in series: the first as above, and the second 6 hours later on a new plate containing diet coated with the second bacterial strain. As with the first exposure, larvae were exposed to bacteria in the second inoculum for 6 hours, after which they were returned to a sterile plate with sterile diet.

To examine competitive interactions within the *Nicrophorus* gut, larvae were inoculated either simultaneously or in series with two of the four species in the following pairings (Strain 1 vs Strain 2): P vs S; M vs S; P vs E; M vs E; P vs M; and S vs E. Within each treatment, larvae from independent families (n = 6) were inoculated as outlined above and then 6 larvae were destructively sampled for plating 6h or 24h later. These values were taken as estimates of input and final densities, respectively. Competition indices (CI) were calculated using the following equation: CI = (Strain 1 output /Strain 2 _output_) / (Strain 1 _input_ / Strain 2 _input_), where input and output values refer to initial and final densities of each competitor, respectively. CI w log transformed, so that a CI of 0 indicates equal competitiveness, while CI > 0 indicates that the strain 1 is a stronger *in vivo* competitor. A two-tailed t-test was used to test if CI values for each strain differed significantly from 0.

At each time point, larvae were sampled by sterilely dissecting individual larval guts with fine forceps and suspending these in 0.7 ml sterile PBS. Gut contents were serially diluted in PBS and plated to quantify CFU on a chromogenic medium (CHROMagar™ Orientation), which can distinguish our experimental strains based on both color and morphology (Figure S1). Mortality rates for each treatment were: PvsS (38.1%); MvsS (47.6%) and PvsM (35%), PvsE (54.5%); MvsE (64.7%) and SvsE (71.4%).

### Larval fitness with different bacterial colonizers

To determine the impact of different bacterial symbionts on larval survival, larvae were inoculated as above and then monitored for survival through time. Larvae exposed to sterile PBS (pH = 7.2) were used as a control in this experiment. A minimum of 40 larvae from 9 ‐ 15 families was collected for each bacterial treatment. We monitored larval survival every 24 hours after inoculation. To reduce the high rates of mortality in larvae reared on liver, all the experimental larvae were transferred daily into a fresh petri dish containing fresh sterile diet.

### Statistical Analysis

Parental fitness and bacterial colonization data were analyzed using ANOVA. Larval survival data was analyzed by fitting a Cox proportional hazard model; this model was constructed by fitting a saturated model using treatment, block and treatment*brood interactions as covariates. The Wald’s test was used to compare mortality between treatments. All analyses were conducted using SPSS version 24 (IBM SPSS Inc., Chicago, IL, U.S.A.).

## Results

### The effects of gut microbiota on parental fitness

To examine the role of the parental gut microbiome on beetle fitness, we reared larvae either with (FC) or without parental care (NC) and then mated the dispersed adults within treatments and allowed them to rear broods on fresh carcasses. Through this treatment, all parents were given the opportunity to rear offspring under identical conditions, and there were neither differences in carcass or maternal weight (both NS). Previous results have shown that parents from these different rearing conditions differ significantly in microbiome composition (18), the FC individuals containing an endogenous symbiont population and the NC individuals a microbial population derived from the soil and the decomposing carcass. Our results show that gut microbiomes have a significant influence on parental fitness (Figure 1). FC parents produced significantly heavier broods than NC parents, irrespective of brood size (2‐tailed ANCOVA: F_1,27_ = 6.09, p = 0.021). In addition, the mean larval mass was heavier in the FC group when controlling for brood size; however, this difference is not significant (2-tailed ANCOVA: F_1,27_ = 3.69, p = 0.067).

**Figure 1.**
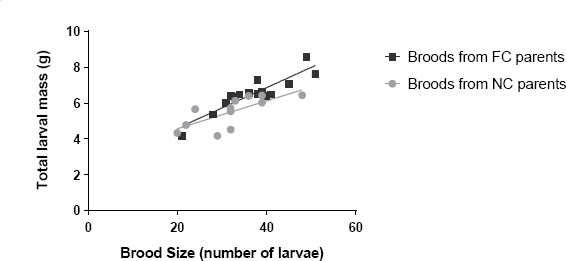
Total larval mass as a function of brood size for maternal beetles that were reared with either Full Care (FC) or No Care (NC).

### Competitive interactions *in vivo*

To study competitive interactions between bacteria during larval colonization, we selected four focal species to examine in detail. Two species, *Providencia rettgeri* and *Morganella morganii* are common endogenous colonizers of the beetle gut (18, 19), while the other two, *S. marcescens* and *E.coli,* are found more commonly in the guts of beetles reared without parental care (18). We first inoculated beetles with each species alone to measure growth and colonization. Figure 2 shows that while all four species are able to colonize the larval gut, their ability to increase in density *in vivo* varies significantly between strains (2-tailed ANOVA: F_3,12_ = 43.13, p < 0.001).

**Figure 2.**
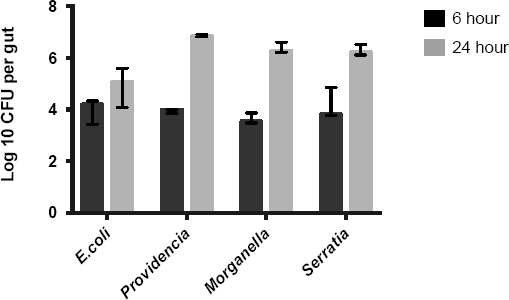
ToGrowth and colonization of *N*. vespilloides symbionts within the larval gut over 24 hrs. Values correspond to the mean +/- 95% CI.

Competitive interactions between species were next determined using *in vivo* pairwise assays where two species were simultaneously inoculated into 1 day old larvae. Consistent with expectations based on mono-associated larvae, we observed clear competitive differences between the strains. *Providencia* and *Morganella* significantly outcompeted both *Serratia* (P vs S: t_5_ = 2.52, p = 0.053; M vs S: t_3_ = 4.42, p = 0.022) and *E. coli* (P vs E: t_2_ = 11.26, p < 0.001; M vs E: t_3_ = 5.89, p = 0.01), although to different degrees. By contrast, there were no significant competitive differences between *Providencia* and *Morganella* (P vs M: t_4_ = 2.16, p = 0.097) (Figure 3A).

**Figure 3.**
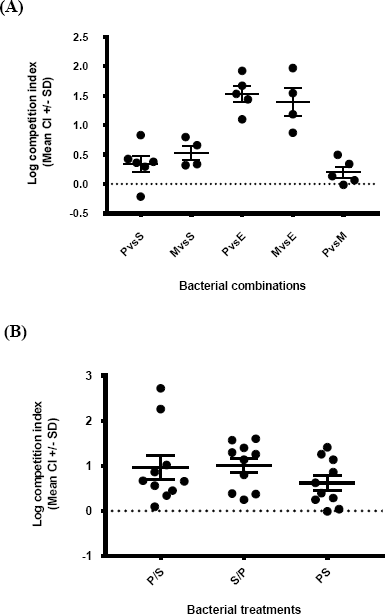
Competitive differences between different bacterial species *in vivo* within the larval gut. Competition indices (CI) are given in reference to the first species listed on the x-axis for (A) and with respect to *Providencia* for (B). Strains were either inoculated simultaneously (A) or in series (B) in cases where strains are separated with a / (e.g. P/S: *Providencia* was inoculated first and then followed with *Serratia,* whereas in PS both strains were co-inoculated). The dashed black line illustrates a CI of 0, which indicates equal competitiveness of two strains. Values > 0 indicates that strain 1 is a stronger *in vivo* competitor.

We next determined if competitive interactions between *Providencia* and *Serratia* were influenced by the order of inoculation. Specifically, we were interested in determining if the outcome of competition was reversed in larvae that were first inoculated with *Serratia.* Our results in Figure 3B clarify that order is not an important determinant of competitive fitness (2-tailed ANOVA: F_2,29_ = 0.59, p =0.56). *Providencia* outcompetes *Serratia* in all cases to a similar degree regardless of the order of inoculation.

### Larval survival with different bacterial colonizers

Our results show that the endogenous microbiome provides likely benefits to *Nicrophorus* by increasing total brood mass, and also that key members of this microbiome can outcompete species that are predominantly found in larvae that do not receive parental care. To test if these competitive interactions translate into differences in larval fitness, we measured the survival of larvae inoculated with single or multiple strains, as above. Results in Figure 4A show that larval mortality varies significantly as a function of their bacterial colonists (χ^2^ = 11.364, df = 3, P < 0.01), with increased mortality in larvae inoculated with *Serratia* compared to either *Morganella, Providencia* or a PBS control (Wald statistic = 6.274; 4.794; 9.202, respectively, all P <0.05). This result is consistent with the known pathogenic effects of *Serratia.* By contrast, there were no significant differences in mortality between larvae inoculated with either *Providencia* or *Morganella* and the PBS control (χ^2^ = 0.156, df = 2, P = 0.925).

**Figure 4.**
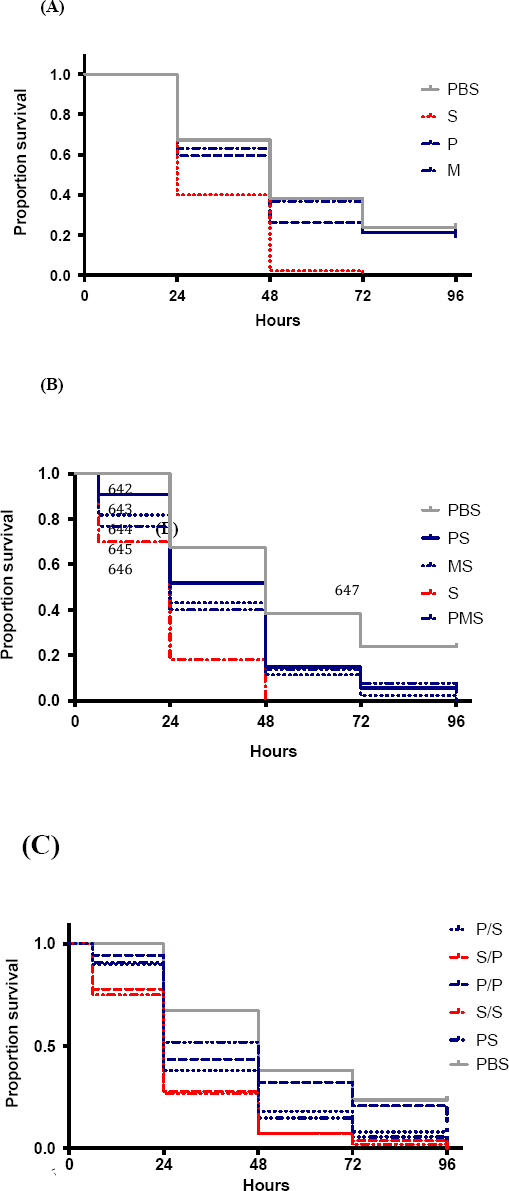
Larval survival when inoculated with different bacterial species. Larvae were inoculated with (A) single bacterial species in monoculture, (B) > 1 species in coculture simultaneously, or (C) bacteria either simultaneously or in series. Bacteria inoculated simultaneously are designated with the first letter of the species name (e.g. PS = *Providencia* with *Serratia),* while species inoculated in series are given in the same way with a slash (e.g. P/S = *Providencia* followed by *Serratia).* A PBS (phosphate buffered saline) control was set up for all the experiments.

We also observed significant differences in survival when larvae were simultaneously inoculated with *Serratia* and either of the endogenous species compared to survival when *Serratia* is inoculated alone (χ^2^ = 38.767, df = 4, P < 0.001). Most importantly, we found that co-inoculating *Serratia* with *Providencia* and/or *Morganella* significantly increased larval survival, suggesting that these species provide protection via colonization resistance for larvae (Wald statistic of PS; MS; PMS = 8.188; 3.697; 5.102, respectively, all P < 0.05, Figure 4B). However, this benefit of colonization resistance disappeared when *Serratia* was able to become established prior to inoculation with *Providencia;* there were no survival differences between larvae inoculated with *Serratia* twice in series and larvae first inoculated with *Serratia* and then followed by *Providencia* (Wald statistic of S/P = 0.077, P = 0.782, Figure 4C). In light of results in Figure 3B showing that order of inoculation does not affect *Providencia* competitiveness, these survival results indicate that *Serratia* induced larval mortality is insensitive to bacterial competitiveness, thus supporting the idea that initial establishment of the endogenous microbiota is crucial for colonization resistance.

## Discussion

*Nicrophorus* larvae are exposed to a highly diverse microbiota in their breeding environment, first from the soil where they hatch and next from the microbes proliferating on and within their carrion resource. In the absence of parental care, larvae become colonized with these bacteria (18) which reduces their weight and survival (15), and also leads to reduced brood mass when these larvae reproduce as parents (Figure 1). However, when larvae are reared with parental care, their gut microbiome resembles that of their parent, even if parental care is limited to carcass preparation prior to larval hatch (18). These results suggested that the bacteria within the parental gut are better competitors for the larval gut, but our earlier work neither tested the colonization potential and competitiveness of the constituent species nor determined the consequences of colonization with the *Nicrophorus* “endogenous” microbiome. Our aims here were therefore to address these questions experimentally by inoculating different endogenous or non-endogenous bacterial species into the guts of developing larvae. We focus specifically on four species: *Providencia rettgeri* and *Morganella morganii,* that are dominant members of the larval microbiome (18, 19), and *Escherichia coli* and *Serratia marcescens,* which are non-endogenous species, but which are either observed in the larval gut *(Serratia)* or have the potential to colonize it through exposure on the mouse carcass (*E. coli*) (18, 32).

Using this approach, we first determined that there are clear differences in the colonization potential of different bacterial species. While *Providencia, Morganella* and *Serratia* increase in density more than 100-fold in 24 hours within the larval gut, *E. coli* was a poor colonizer and only increased by ∼ 10-fold over the same time interval (Figure 2). In addition to clarifying these differences, these experiments also established that it is feasible to experimentally colonize larval beetles via diet manipulation. The growth differences between strains in monoculture were reflected in their interactions *in vivo* during co-culture. Specifically, we saw competitive dominance of *Providencia* and *Morganella* over *E. coli* and *Serratia* when pairs of strains were simultaneously fed to larvae (Figure 3A). Moreover, in competition experiments between *Providencia* and *Serratia,* we found that the order of inoculation did not affect the competitive outcome between strains (Figure 3B). This latter result suggested that priority effects are not realized in this system because *Serratia* could be displaced even after a 24-hr head start in colonization. By contrast, another recent study found that the colonization competitiveness of *Borrelia* strains within the mouse gut are significantly determined by their order of presentation to the host mouse (33).

At present, we have limited understanding of the factors that mediate the competitive differences between strains within the *Nicrophorus* larval gut. Differences in *in vivo* growth rates are sufficient to explain the competition results during simultaneous inoculation. However, the fact that *Providencia* can still invade an established Serratia-colonized larva (Figure 3B), suggests the possibility that competitive interactions are in part mediated by the host. For instance, host innate immunity could be a direct factor in determining the competitive outcome and final population density of bacterial species within hosts (34). Equally, commensal bacteria could prime the host immune response to limit pathogen colonization by causing an up-regulation of antimicrobial peptides, such as AMP molecules in *Aedes aegypti* mosquitoes and islet-derived protein 3γ in mice (35, 36); however, these would need to be specifically targeted to non-symbiont species. Host involvement in this system is further suggested by other experimental results showing that *in vitro, Serratia* is able to outcompete *Providencia* (YW unpublished data), a result likely attributed to the faster growth of this strain during *in vitro* culture. An important aim for future work will be to clarify the factors that drive the competitive interactions between bacterial strains during colonization.

To understand the consequences of the *Nicrophorus* microbiome for beetle fitness, we quantified larval mortality following inoculation of monocultures or co-cultures of different bacterial species. Consistent with results on colonization resistance in other systems (9, 37), these experiments showed that *P. rettgeri* and *M. morganii* both provide protection against *Serratia* infection, but with no added protection if both endogenous species are present (Figure 4B). This is expected given the results from *in vivo* competition assays. By contrast, when *Serratia* is inoculated first, the protection provided by *Providencia* is abolished, in spite of the fact that *Providencia* can outcompete *Serratia* in these conditions (Figure 4C). Interestingly, these results indicate that the pathogenesis of *Serratia* is separate from its *in vivo* competitive ability, perhaps owing to toxin production or invasion through the gut into the haemocoel within the first 24 hours (38–40). Thus initial establishment of the endogenous microbiota is apparently crucial for colonization resistance.

Although the mechanism of *Serratia*-induced mortality remain unknown in this species, the fact that colonization resistance requires the prior or simultaneous establishment of the *Nicrophorus* endogenous microbiota has important implications for our understanding of the functions of parental care. Parents protect larvae and provide nutrition in the form of regurgitated food (22). In addition, they transfer their gut microbiome to larvae by direct feeding and via contamination of the carcass surface (17, 18). The present results indicate that larvae benefit directly from the acquisition of these bacteria (Figures 1 and 4B), and suggest that the microbiome, or at least two of its key members, are mutualists with *Nicrophorus.* Thus while *Nicrophorus* adults ensure transmission of these species from generation to generation, the bacteria provide direct benefits to beetles within the highly contaminated carcass environment (19). It remains possible that other advantages exist, for example improved nutrient acquisition (41) or changes in the composition of the decomposer microbial community on the carcass (17), but as yet these possibilities have not been measured.

Our results provide strong evidence that members of the *Nicrophorus* microbiome provide direct advantages to larvae and adults; however, it is important to note that these advantages were not measured in the natural context of the carcass itself. While this was necessary for the current work, it does mean that we may be underestimating larval exposure to potential bacterial pathogens (42). In addition, our assay clearly suffers from extremely high rates of larval mortality, irrespective of treatment. Artificial diets that better mimic the larval environment and that improve larval nutrition and survival are thus needed to more fully elucidate the functions of the *Nicrophorus* microbiome. However, despite these limitations, our results point towards yet another role of parental care in *Nicrophorus vespilloides,* and argue for further comparative studies in other congeners that vary in their requirements for parental care.

## Acknowledgements

We acknowledge the helpful comments of Chris Jacobs and Andres Arce on an earlier version of this manuscript. YW was supported by a graduate scholarship from the China Scholarship Council and DER was supported by funds from Leiden University.

---

**Figure S1.**
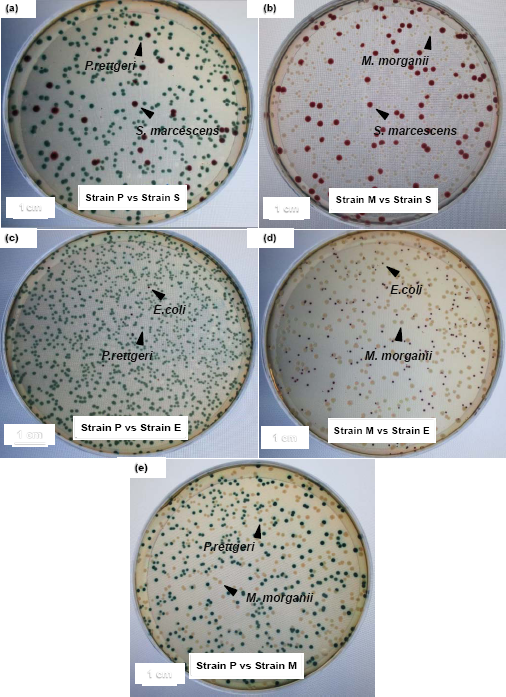
Color and morphology of experimental strains on chromogenic agar plates (CHROMagar™ Orientation) used for bacterial competition assays. Bacterial combinations shown are: P vs S (a); M vs S (b); P vs E (c); M vs E (d); P and M (e).

